# Functional specialization and dynamical interaction in human amygdala subregions support fearful-expression recognition

**DOI:** 10.1101/2025.11.16.688165

**Authors:** Dan Cao, Jiachen Cui, Xinyuan Yan, Xinru Zhang, Yuanyuan Dang, Hulin Zhao, Jin Li, Jianning Zhang, Yanyang Zhang

## Abstract

Fearful-expression recognition is critical for adaptive responses to potential threats and relies on both rapid threat detection and fine-grained face encoding. Yet how human amygdala subregions differentially support these distinct cognitive components remains unclear. Here, we recorded intracranial EEG from lateral and medial amygdala in epilepsy patients performing an emotional face-matching task and combined multivariate decoding, time-frequency and directed-connectivity analyses with intracranial stimulation. The lateral amygdala exhibited early fear-specific responses, characterized by higher decoding accuracy and increased theta/alpha-band (4-12 Hz) power, and transmitted this fear-related information to the medial amygdala, which showed delayed and sustained activation. By contrast, the medial amygdala encoded face-specific information at later stages in the 2-16 Hz band with superior decoding accuracy and then relayed this information back to the lateral amygdala. Intracranial stimulation produced a double dissociation in behavior, with lateral amygdala stimulation disrupting fear detection, whereas medial amygdala stimulation selectively accelerated neutral-face recognition. Together, these findings reveal a temporal hierarchy in the human amygdala, whereby dynamic bidirectional interactions between subregions implement distinct components of fearful-expression recognition, providing a circuit-level framework for understanding social threat processing.

## Introduction

Recognizing fearful expressions is a core survival function, rapidly directing attention to potential threats and mobilizing physiological and behavioral responses ^1^. The amygdala is a principal subcortical hub for this process ^2^. However, successful threat evaluation requires not only rapid detection of fear cues but also fine-grained encoding of facial features, and these complementary components jointly shape adaptive emotional and behavioral outcomes ^1,3^. The amygdala’s rapid sensitivity to fear is well documented ^4,5^. Recent studies further report that amygdala neurons can encode non-emotional facial information, responding selectively to face parts such as the eyes and mouth ^6–8^. These evidence characterize the amygdala as a processor of multidimensional information ^9^. Yet many studies treat the amygdala as a unitary structure despite established anatomical and functional heterogeneity across its subregions ^10,11^. Accordingly, how the amygdala and its subregions contribute to distinct cognitive components of fearful-expression recognition remains unclear.

Rodent studies have demonstrated that the amygdala comprises anatomically and functionally heterogeneous nuclei. The basolateral complex (BLA) receives extensive cortical inputs and is implicated in affective evaluation, whereas the central nucleus (CeA) projects to hypothalamic and brainstem targets to drive autonomic and behavioral outputs ^12^. In humans, studies parcellated the amygdala into lateral and medial subdivisions that broadly map onto BLA-like and CeA-like regions ^10,13^. Structurally, diffusion-based tractography shows that these lateral and medial subregions exhibit distinct white-matter connectivity profiles with cortical and subcortical targets, supporting their anatomical differentiation ^14,15^. Functionally, connectivity-based fMRI and intracranial stimulation studies reveal separable whole-brain coupling and spatiotemporal activation patterns for lateral and medial sites ^16,17^. Moreover, monosynaptic rodent tracing and human lesion studies point to inter-subregional interactions, indicating that lateral-medial communication may be important for emotional responses ^18,19^. Collectively, these lines of evidence suggest a division and cooperation between the two subregions. This raises a critical question: whether and how the amygdala subregions dominate and interact to support distinct processes of fear detection and face encoding.

Preliminary human evidence indicates a functional dissociation between lateral and medial amygdala subregions during fear and face processing. Lesion studies show that patients with BLA damage exhibited impaired fear recognition ^20^, deficient fear memory ^21^ and increased generalization of fear responses ^22^. Complementary single-unit recordings reveal that neurons in the medial subdivision can carry higher-order face representations (e.g., identity, feature-selective codes) ^7^. Nevertheless, direct comparisons of lateral versus medial subregions during fear or face processing remain scarce and have yielded mixed results. For example, some fMRI studies have reported stronger medial activation during fear processing ^23,24^, whereas others have found greater lateral responses ^25,26^. Although these studies provide valuable insights, they are constrained by the indirect and temporally coarse nature of the BOLD signal, which limits their ability to resolve rapid neural dynamics and directed interactions between subregions that are critical for understanding threat and face processing.

To address these issues, we recorded intracranial EEG (iEEG) activity simultaneously from lateral and medial amygdala sites in 12 drug-resistant epilepsy patients while performing an emotional face-matching task (EFMT) ^27^ (**Fig. 1A**). By integrating time-resolved decoding, spectral power (**Fig. 1B**), information directionality (**Fig. 1C**) analyses, and intracranial electrical stimulation (iES; **Fig. 1D**), we investigated how lateral (lAmyg) and medial (mAmyg) amygdala subregions dynamically specialize and interact to support fearful-expression recognition. We found that lAmyg was engaged early in fear-specific processing, exhibiting higher decoding accuracy and elevated theta/alpha-band power, and transmitted this information to mAmyg. Conversely, mAmyg dominated late-stage face processing, with superior decoding accuracy and 2-16 Hz spectral power, and then relayed face-specific information back to lAmyg. Moreover, iES in seven additional patients showed that stimulating lAmyg or mAmyg selectively modulated behavioral responses to fear detection and face recognition, respectively, providing causal evidence for their distinct functional contributions.

**Fig. 1.** Overall study framework. (A) Participants completed an emotion face matching task, with intracranial EEG (iEEG) recorded from lateral (lAmyg) and medial (mAmyg) amygdala subregions. (B) Local activity analyses: lAmyg showed early, fear-specific responses with higher time-resolved decoding accuracy and elevated theta/alpha-band power (pink panel); mAmyg encoded late, face-specific information in the 2-16 Hz band with superior decoding accuracy and increased spectral power (light blue panel). (C) Information directionality analyses: lAmyg sent fear-specific information to mAmyg early, while mAmyg transmitted face-specific information back to lAmyg later. (D) Intracranial stimulation: causal perturbations of lAmyg and mAmyg yielded distinct behavioral modulations.

## Results

### Task, behavior and recording channels

Twelve patients with drug-resistant epilepsy (3 females, **Table 1**) completed an EFMT task during invasive pre-surgical evaluation. The task comprised three sequential blocks corresponding to the fearful-face, neutral-face, and control-shape conditions.

**Table 1.** Participants characteristics with iEEG recordings.

| Patient | Age | Sex | lAmyg sites<br>(response) | mAmyg sites<br>(response) | Seizure onset zone |
| --- | --- | --- | --- | --- | --- |
| 1 | 21 | male | 3(3) | 3(3) | Hippocampus, right |
| 2 | 31 | male | 1(1) | 0 | Hippocampus, left |
| 3 | 16 | male | 2(1) | 2(2) | Hippocampus, right |
| 4 | 47 | male | 3(3) | 3(3) | Parietal lobe, right |
| 5 | 22 | male | 3(2) | 0 | None |
| 6 | 32 | male | 5(5) | 4(4) | Cingulate gyrus, right |
| 7 | 50 | female | 6(6) | 0 | None |
| 8 | 27 | female | 2(2) | 0 | Hippocampus, left |
| 9 | 28 | male | 1(1) | 0 | Hippocampus, right |
| 10 | 54 | female | 5(5) | 2(2) | None |
| 11 | 31 | male | 1(1) | 3(2) | Hippocampus, left |
| 12 | 24 | male | 6(1) | 3(1) | None |

Abbreviations: lAmyg, lateral amygdala; mAmyg, medial amygdala

In each block, the participants were instructed to match one of two images presented at the bottom of the screen to a target image displayed at the top within a 4-second window (**Fig. 2A**, see Methods for details). We found that the recognition accuracy for the fearful condition was the lowest (mean ± SD: 68.75% ± 21.28%), followed by that for the neutral faces (mean ± SD: 85.07% ± 22.99%), while the accuracy in the shape condition was the highest (mean ± SD: 90.97% ± 14.84%) (repeated-measures analysis of variance [ANOVA]: *F*(2, 22) = 8.67, *p =* 0.001; **Fig. 2B**). Moreover, the participants exhibited significantly longer reaction times to the fearful faces (mean ± SD: 2.66 ± 0.35 sec), with those to the neutral faces being intermediate (mean ± SD: 2.04 ± 0.62 sec) and to the control shapes being the shortest (mean ± SD: 1.34 ± 0.38 sec) (ANOVA: *F*(2,22) = 55.77, *p* < 0.001; **Fig. 2C**). These behavioral findings indicate that the recognition of fearful faces is more difficult, which is consistent with previous study that fearful expressions requires greater cognitive resources for processing ^28^.

**Fig. 2.**
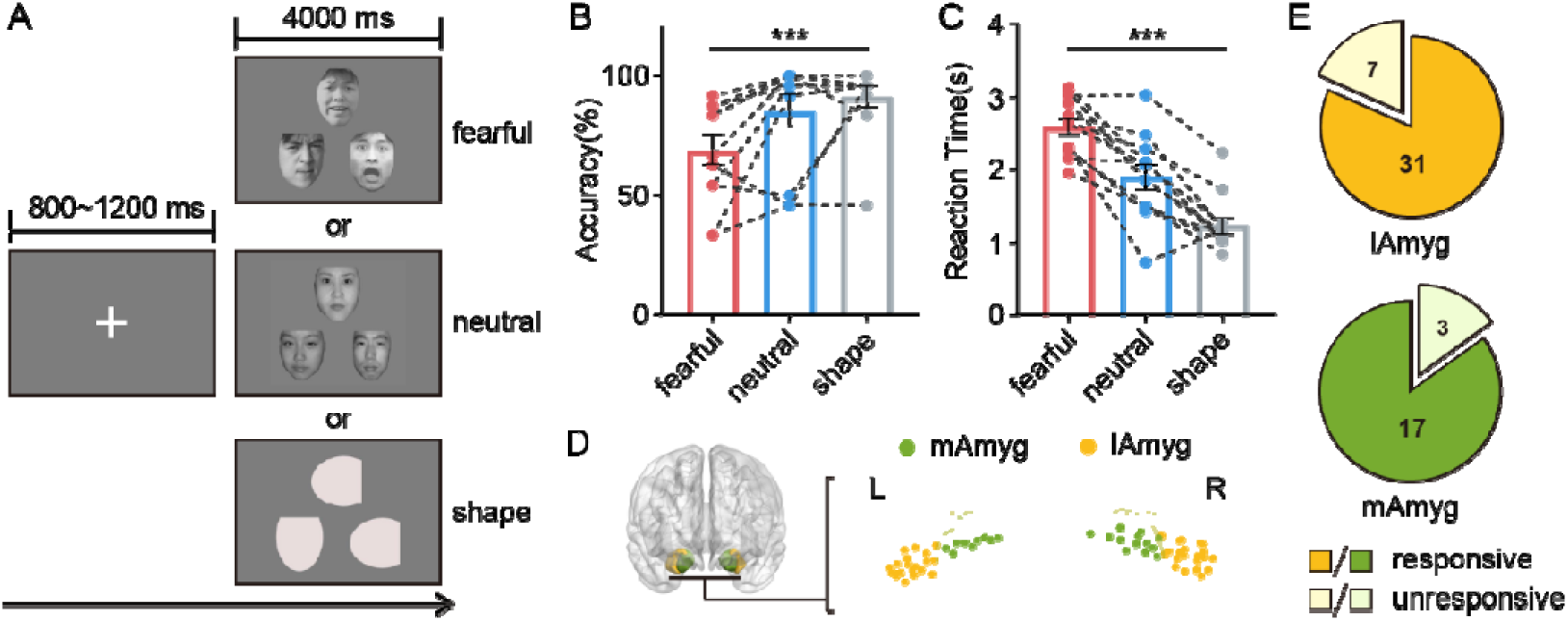
Experimental task, behavioral result and recording contacts. (A) An example trial of the task, showing example stimuli for each condition (fearful, neutral and shape). The stimulus was presented for 4 seconds, and participants were required to judge which of the two images below matched the expression or property of the image above during this period. (B) Accuracy differed across conditions, with the lowest for fearful faces and the highest for shape (*F*(2, 22) = 8.67, *p* < 0.001). Bar graphs indicate the mean accuracy per condition. Each dot represents each participant and black dashed lines connect dots from the same participant across conditions. *** *p* < 0.001. (C) Reaction times were different across conditions. Bar graphs indicate the mean reaction time, with the longest for fearful faces and the shortest for shape (*F*(2, 22) = 55.77, *p* < 0.001). Each dot represents each participant and black dashed lines connect dots from the same participant across conditions. *** *p* < 0.001. (D) Electrode location across participants in Montreal Neurologic Institute (MNI152) space. Recording subregions are indicated by different colors(orange, lAmyg; green, mAmyg). The brain figure was visualized by BrainNet Viewer toolbox (www.nitrc.org/projects/bnv/) ^30^. (E) Responsive contacts in the mAmyg (81.6%, 17/20) and the lAmyg (85.0%, 31/38).

Local field potentials (LFPs) were recorded from depth electrodes implanted in medial (mAmyg, 20 contacts) and lateral amygdala (lAmyg, 38 contacts; **Fig. 2D**) across participants. To ensure that the subsequent analyses targeted stimulus-related responses, responsive contacts were selected as described in previous study ^29^. Specifically, the contact exhibiting *z*-scored power values exceeding 1.96 for at least 50 ms was defined as a responsive contact (see Methods for details). For each condition, we found that the largest proportion of responsive contacts across all frequency bands (1-150 Hz) occurred under the fearful condition (lAmyg: 91.22% ± 8.02%, mAmyg: 95.79% ± 6.86%; **Fig. S1B**), followed by the neutral condition (lAmyg: 77.56% ± 14.05%, mAmyg: 82.11% ± 15.16%), and the lowest in the shape condition (lAmyg: 73.17% ± 13.90%, mAmyg: 72.63% ± 21.82%).

Therefore, we included those contacts that exhibited full-band responses under at least one condition. As illustrated in **Fig. 2E**, 81.6% of the contacts in lAmyg (31/38) and 85.0% of the contacts in mAmyg (17/20) were responsive. These responsive contacts were subsequently included in the analyses.

### Temporal dissociation of amygdala subregional engagement in fear-specific processing

Given that the participants consistently responded within 3 s following stimulus onset (**Fig. 2C**), subsequent analyses were restricted to the 0-3 s window to avoid contamination from unrelated neural activity to the task. For each amygdala subregion, we computed time-frequency power during stimulus presentation and *z*-scored these values relative to pretrial baseline distributions to quantify task-induced power changes at the single-trial level. Responsive contacts within both subregions exhibited sustained low-frequency activity (2-16 Hz, *z* > 1.96, *p* < 0.05; **Fig. 3A**) across both the fearful and neutral conditions. These findings mirrored the time-frequency power observed across all contacts within the two subregions (**Fig. S1A**), suggesting that the selected responsive contacts captured the essential activation patterns of the amygdala. To test the fear-specific effect, we compared the *z*-scored power of both subregions between the fearful and neutral conditions using a cluster-based permutation test. This analysis revealed greater theta/alpha-band (4-12 Hz) power for the fearful condition than the neutral condition within both lAmyg (*p* < 0.001; **Fig. 3A** top) and mAmyg (*p* < 0.001; **Fig. 3A** bottom).

**Fig. 3.**
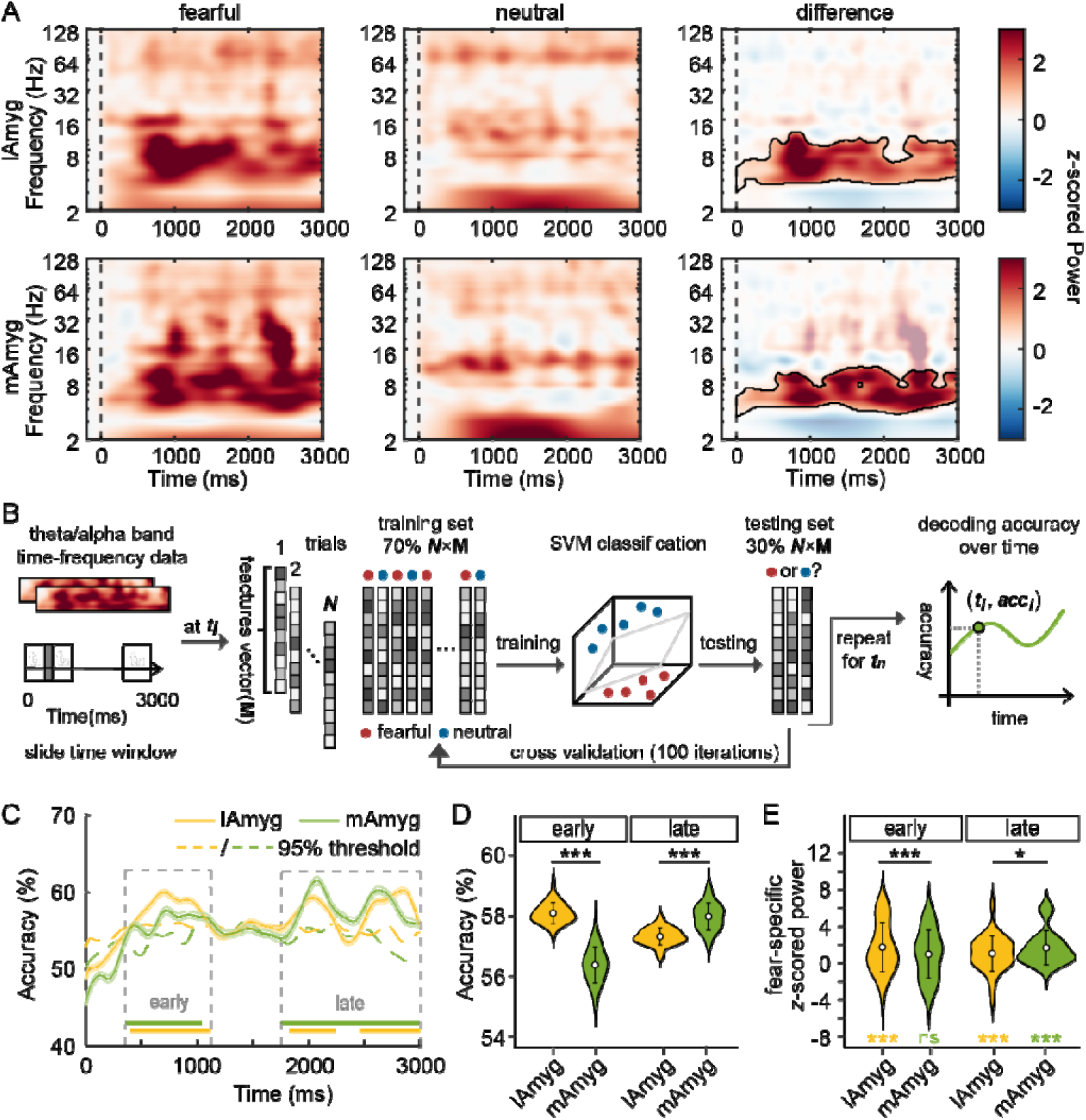
Temporal dissociation of amygdala subregions during fear-specific processing. (A) Time-frequency representations for the fearful (left), neutral (middle) and their contrast (fearful - neutral) conditions, separately for the lAmyg (top) and mAmyg (bottom). Compared to the neutral condition, the fear condition elicited significantly increased power in the theta/alpha band (4-12 Hz), as highlighted by the black lines (all *ps* < 0.05). (B) The schematic of the time-resolved decoding analysis. For each time window *t*, the theta/alpha power features were extracted from all fearful and neutral trials for each subregion. The SVM classifier was trained on 70% of trials and tested on the remaining 30%, with 100 repeated random 70/30 splits; repeating this across all windows produced a time-resolved decoding accuracy curve per subregion. (C) Time-resolved decoding accuracy for the two subregions. Shaded areas represent standard error of the mean, the dashed line indicates the 95^th^ threshold derived from the random distribution, and the bottom horizontal bars mark time intervals where decoding accuracy exceeded the threshold. Gray dashed boxes indicate the early (360-1110 ms) and late (1750-3000 ms) windows identified from the significant decoding accuracy (all *ps* < 0.05). (D) Mean decoding accuracy in the early and late time windows for each subregion. The lAmyg exhibited significantly higher decoding accuracy in the early window, while the mAmyg showed superior decoding accuracy in the late window. White points in violins represent the mean value; vertical black lines indicate the standard deviations. *** *p* < 0.001. (E) Fear-specific theta/alpha power (fearful - neutral) in the defined early and late time windows, with white points and black lines represent the mean value and standard deviations. The lAmyg showed larger fear-specific power increase in the early window(*t* = 3.52, *p* < 0.001), while the mAmyg exhibited greater fear-specific power enhancement in the late window (*t* = -2.15, *p* = 0.037). * *p* < 0.05; *** *p* < 0.001.

Further, to investigate the temporal dynamics of each subregion during the fear-specific processing, we conducted a time-resolved multivariate decoding analysis using the theta/alpha band power as input features, since multivariate decoding approaches can capture information that is lost when averaging signals, providing greater sensitivity for detecting differences between conditions ^31^. **Fig. 3B** shows the schematic of the time-resolved decoding analysis. We extracted power features from lAmyg and mAmyg, then used a sliding-window linear support vector machine (SVM) classifier to decode fearful vs. neutral stimuli, yielding decoding accuracy over time. Statistical significance was assessed via a null distribution generated by shuffling label-data relationships (see **Methods** for details). Results indicated a temporal reversal in decoding accuracy, with lAmyg showing higher accuracy in the early window while mAmyg showing superior in the late window. Specifically, both subregions exhibited significant decoding accuracy (exceeding the 95th percentile of the null distribution, all *ps* < 0.05; **Fig. 3C**) across two temporal clusters: for lAmyg, these were 400-1100 ms and 1830-3000 ms; for mAmyg, 360-1040 ms and 1750-3000 ms. We then defined an early window (360-1110 ms) and a late window (1750-3000 ms) based on above clusters and extracted the mean accuracy for each subregion in these windows. In the early window, lAmyg outperformed mAmyg (58.1% ± 0.3% vs. 56.4% ± 0.6%; paired t-test: *t* = 25.10, *p* < 0.001; **Fig. 3D**). In contrast, mAmyg surpassed lAmyg in the late window (58.0% ± 0.4% vs. 57.3% ± 0.3%; *t* = 12.87, *p* < 0.001).

Next, to assess whether the fear-specific power differed between subregions in the early and late windows, we defined the fear-specific power as the theta/alpha *z*-scored power in the fearful condition minus that in the neutral condition. We then compared the fear-specific power between the two subregions for each window using a linear mixed-effects model, with subregion as the fixed variable, fear-specific power as the dependent variable, and contact within subject as the random variable. As indicated in **Fig. 3E**, in the early window the lAmyg exhibited a fear-specific power increase (fearful > neutral; linear mixed-effects model: *t* = 4.44, *p* < 0.001), which was significantly larger than in the mAmyg (linear mixed-effects model: *t* = 3.52, *p* < 0.001). While the mAmyg did not show an early enhancement (*p* = 0.063). In the late window both lAmyg (*t* = 3.69, *p* < 0.001) and mAmyg (*t* = 4.64, *p* < 0.001) showed fear-specific increases, but the mAmyg increase exceeded that of the lAmyg (linear mixed-effects model: *t* = 2.15, *p* = 0.037). Collectively, these findings reveal a temporal dissociation in the fear-specific engagement across amygdala subregions, with lateral subregion predominance in early processing and medial subregion activation during later stage.

### Lateral amygdala subregion drives early-stage information flow during fear processing

Fear-induced increases in spectral power were found within the theta/alpha band for both the lAmyg and the mAmyg, where early lAmyg dominance and late mAmyg dominance were identified in fear processing within this band. This raised a further question about whether the two subregions of the amygdala interact during fear processing. We hypothesized that the lAmyg transfers information to the mAmyg during fear processing. To test this, we first examined the functional connectivity between these subregions via debiased weighted phase lag index (dwPLI). This approach quantifies inter-regional phase synchronization while minimizing volume-conduction artifacts ^32^. Specifically, the time-frequency dwPLI was computed for trials with both fearful and neutral conditions and subsequently *z*-scored against a shuffle-based null distribution (see **Methods**). As shown in **Fig. 4A**, both conditions exhibited significant *z*-scored dwPLI within the low-frequency range (*z* > 1.96, *p* < 0.05), with a between-condition difference found within the theta/alpha band (cluster-based permutation test: *p* = 0.048; **Fig. S2**). Next, we extracted the theta/alpha-band dwPLI and assessed fear-specific connectivity across time. This analysis revealed a significant fear-induced increase in the 691-910 ms time window (cluster-based permutation test: *p* = 0.018; **Fig. 4B**), which falls within the above defined early window. To further examine the stability of this fear-specific effect across individuals, we extracted the fear-specific dwPLI (fearful - neutral) across all contact pairs in the 691-910 ms and found that 35/40 pairs exhibited positive fear-specific dwPLI (linear mixed-effects model: *t* = 4.01, *p* < 0.001; **Fig. 4C**).

**Fig. 4.**
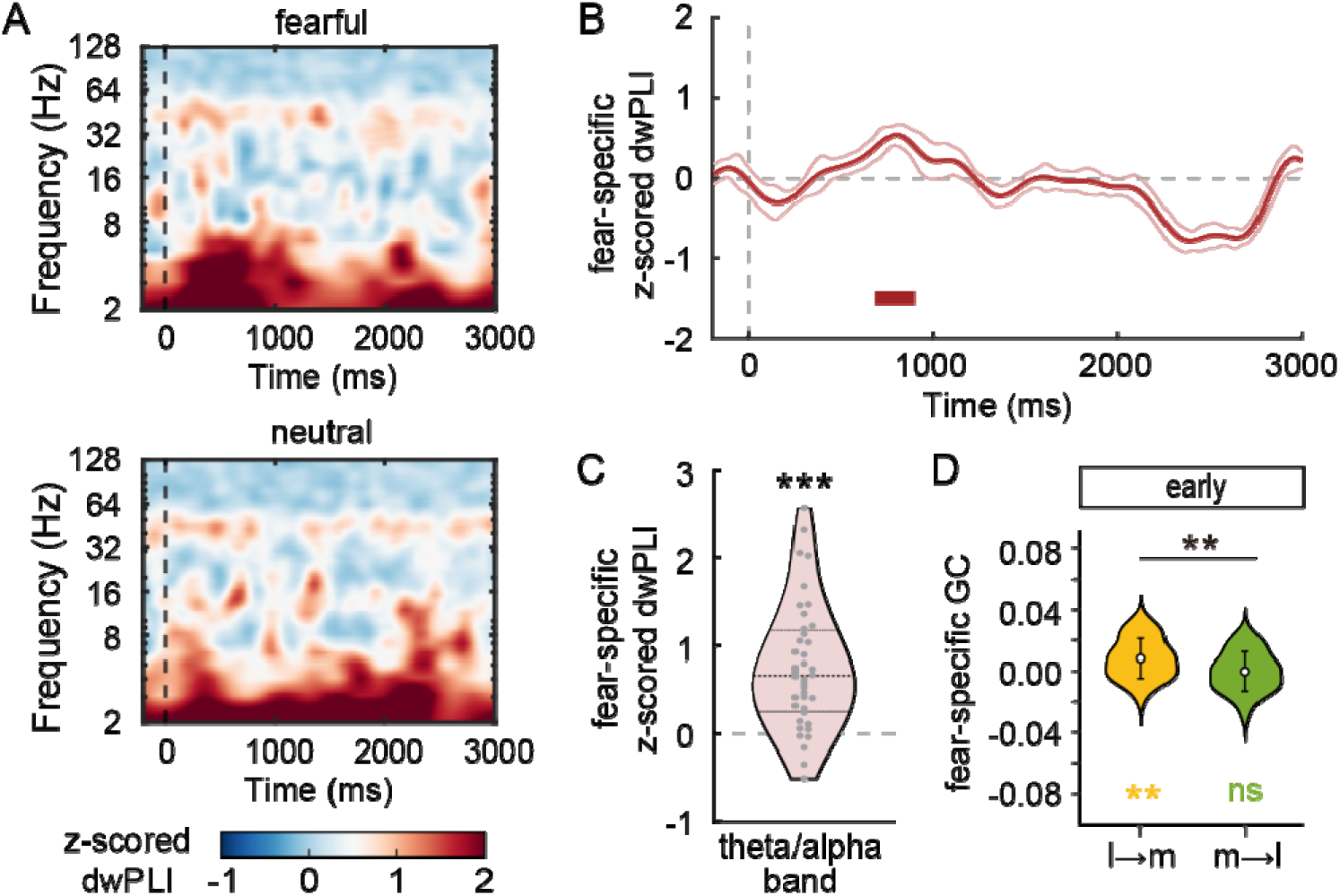
Functional and directional connectivity between amygdala subregions during fear processing. (A) Group-level *z*-scored dwPLI between subregions under the fearful (left) and neutral (right) conditions. Significant phase synchronization was found in the low-frequency range (*z* > 1.96, *p* <0.05). (B) Time course of fear-specific (fearful - neutral) *z*-scored dwPLI with shaded areas representing the standard error of the mean. Red rectangle indicate the time window (691-910 ms) that fear-specific dwPLI enhancement (*p* = 0.018), and gray shaded regions denote the predefined early (360-1110 ms) and late time (1750-3000 ms) windows. (C) Distribution of fear-specific *z*-scored dwPLI across all contact pairs within the 691-910 ms, with black dashed lines indicate the median and interquartile range. Each dot represents one contact pair and a total of 35/40 contact pairs exhibited positive differences (*t* = 4.01, *p* < 0.001). *** *p* < 0.001. (D) Fear-specific theta/alpha-band GC index between the two directions within the predefined early window. White points in violins represent mean values and black lines indicate the standard deviations. In the early window, the GC from lAmyg to mAmyg was fear-specific increased (*t* = 2.67, *p* = 0.009), with a directional dominance over the reverse direction (*t* = 2.94, *p* = 0.004). ** *p* <0.01.

Given that we identified differences in theta/alpha-band dwPLI between the fearful and neutral conditions within the predefined early window, we next employed a non-parametric frequency-domain conditional Granger causality (GC) analysis. This analysis aimed to determine the directionality of fear-specific information flow between the amygdala subregions within the early window. Specifically, we computed the theta/alpha-band GC index with fearful and neutral trials for both lateral→medial (l→m) and medial→lateral (m→l) directions within the predefined early window. And the fear-specific GC was quantified as the difference between the fearful and neutral conditions. Results exhibited that within the early window the theta/alpha-band GC from lAmyg to mAmyg was increased in the fearful condition compared to the neutral condition (linear mixed-effects model: *t* = 2.67, *p* = 0.009), and this increase was greater than in the reverse m→l direction (linear mixed-effects model: *t* = 2.94, *p* = 0.004; **Fig. 4D**). No fear-specific increase was observed for the m→l direction (*p* = 0.747). These results indicated that fear selectively amplifies directed information flow from the lateral to the medial amygdala at early stage, with no analogous effect in the reverse direction or at later stage. Collectively, our findings outline a twolstage fearlprocessing cascade: the lateral amygdala quickly detects and encodes fear early on, then directs this information to the medial amygdala, which becomes more active at late stage.

### Medial amygdala selectively supports late-stage encoding of facial information

Recent studies indicate that the amygdala is not only a hub for emotional evaluation but also participates more broadly in multidimensional information processing ^9^ (e.g., face identity ^7^). Accordingly, we assessed amygdala subregional contributions to face processing using the same pipeline as for fear-specific analyses. We contrasted the neutral-face and control-shape conditions and evaluated functional segregation and interaction among amygdala subregions.

Time-frequency analyses revealed that the neutral faces elicited stronger *z*-scored power than control shapes mainly within the 2-16 Hz range for both the lAmyg and mAmyg (cluster-based permutation test, all *p*s < 0.05; **Fig. 5A**). We then conducted time-resolved multivariate decoding analysis within each subregion using 2-16 Hz z-scored power as features to distinguish neutral-face from control-shape trials. We found that the decoding accuracy of the mAmyg exceeded the 95^th^ percentile of the null distribution in two sustained windows (1090-2340 ms and 2710-3000 ms; all *p*s < 0.05; **Fig. 5B**). By contrast, the lAmyg showed only a brief effect at 1470-1660 ms (*p* < 0.05). Thus, reliable face decoding was mainly observed in the mAmyg during the 1-3 sec post-stimulus period. To further confirm this dissociation, we extracted the mean decoding accuracy from both subregions over the 1090-3000 ms time window and performed a direct comparison. Results showed that the mAmyg exhibited a significantly higher mean decoding accuracy (58.0% ± 1.4%) than the lAmyg (54.0% ± 1.9%; paired t-test: *t* = 23.89, *p* < 0.001; **Fig. 5C**). Moreover, analysis of 2-16 Hz *z*-scored power showed that only the mAmyg exhibited significant face-specific enhancement (face > shape; linear mixed-effects model: *t* = 4.50, *p* < 0.001) within this window (1090-3000 ms), which was also greater than that within the lAmyg (*t* = 3.05, *p* = 0.004; **Fig. 5D**). These results collectively demonstrated that the medial, rather than lateral, amygdala selectively contributes to the face processing in the 1-3 sec post-stimulus interval.

**Fig. 5.**
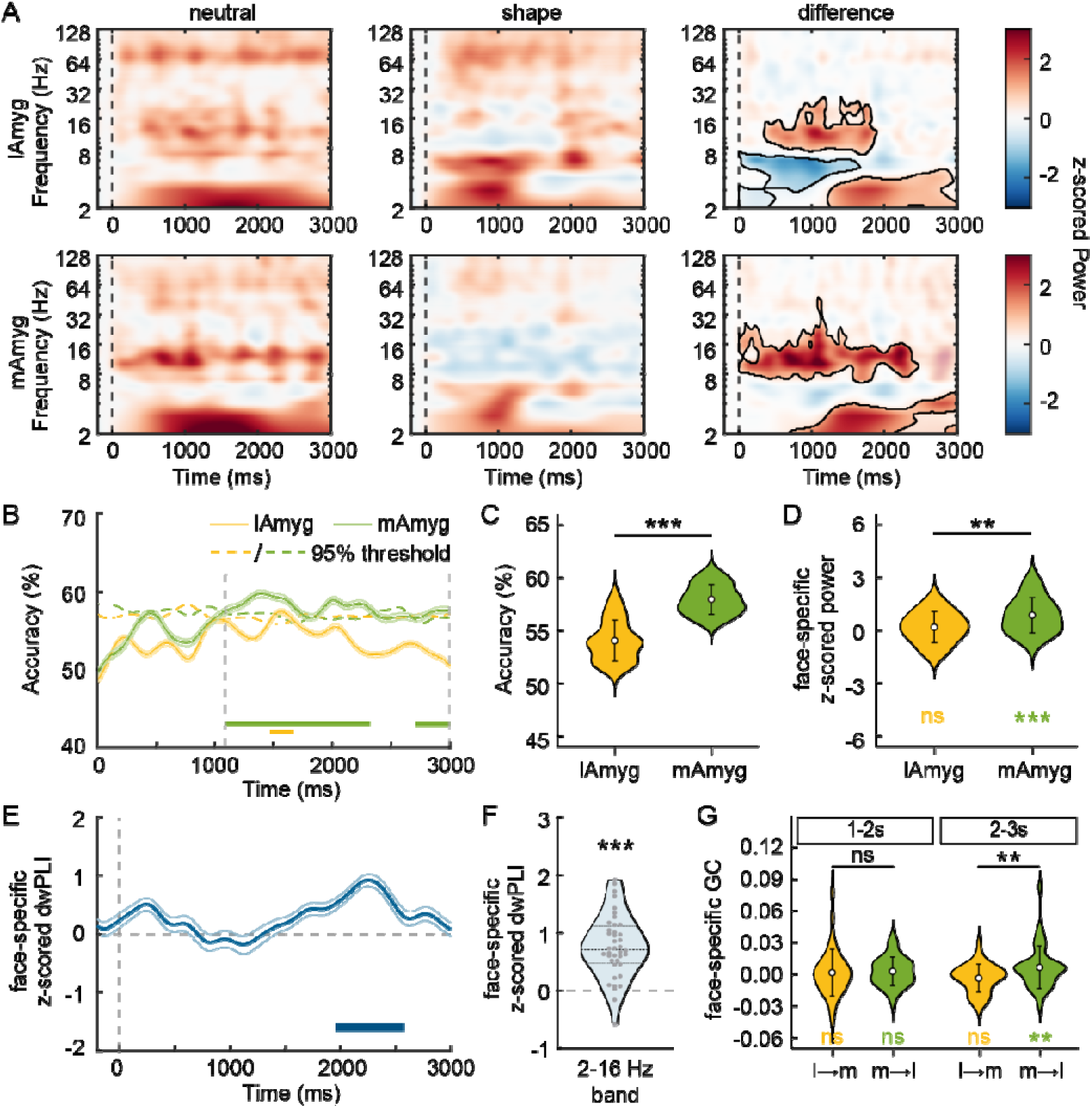
Temporal dynamics of z-scored power and interaction within amygdala subregions during face processing. (A) Time-frequency representations for the neutral condition (left), shape condition (middle) and their contrast (neutral - shape) separately for the lAmyg (top) and mAmyg (bottom). The differences between the two conditions were found within the low-frequency range (2-16 Hz), as highlighted by the black lines (all *ps* < 0.05). (B) Time-resolved decoding accuracy for the mAmyg (green) and lAmyg (orange) subregions, using the 2-16 Hz *z*-scored power features. Shaded areas indicate the standard error of the mean, the dashed line marks the 95^th^ threshold derived from the null distribution, and the bottom horizontal bars mark time intervals where decoding accuracy exceeded the threshold. Gray dashed boxes indicate the time window (1090-3000ms) identified from the decoding results (all *ps* < 0.05). (C) Mean decoding accuracy within significant time window for each subregion. The mAmyg exhibited higher accuracy than the lAmyg. White points in violins represent the mean decoding accuracy; vertical black lines indicate the corresponding standard deviations. *** *p* < 0.001. (D) Face-specific *z*-scored power (neutral - shape) within the 2-16 Hz band over 1090-3000 ms, with white points and black lines represent the mean value and standard deviations. The mAmyg showed larger face-specific power enhancement (*t* = 3.05, *p* = 0.004). ** *p* < 0.01. (E) Time course of face-specific (neutral - shape) *z*-scored dwPLI within the 2-16 Hz range, with shaded areas representing the standard error of the mean. Blue rectangle indicates the time window (1950-2580ms) that face-specific dwPLI enhancement (*p* < 0.001), and gray shaded regions denote the time window (1090-3000ms) by decoding analysis. (F) Distribution of *z*-scored dwPLI difference across all contact pairs within the 1950-2580 ms window, with black dashed lines indicate the median and interquartile range. Each dot represents one contact pair and a total of 38/40 contact pairs exhibited positive differences (*t* = 4.32, p < 0.001). *** *p* < 0.001. (G) Face-specific GC index between the two directions within the 2-16 Hz band. White points in violins represent mean values and black lines indicate the standard deviations. Between 2-3 sec window, the GC index from lAmyg to mAmyg was stronger under the neutral condition (*t* = 2.09, *p* = 0.040), with a clear directional dominance over the reverse pathway (*t* = 2.89, *p* = 0.005). No findings were found in the 1-2 sec window. ** *p* <0.01.

Next, we computed the *z*-scored dwPLI between the two subregions separately for neutral-face and control shape trials, then compared them using a cluster-based permutation test. This comparison revealed a between-condition difference in the 2-16 Hz frequency band (**Fig. S2**). Subsequently, we extracted *z*-scored dwPLI from both conditions within this band and conducted a direct time-resolved comparison. Results showed that neutral faces elicited significantly greater *z*-scored dwPLI than control shapes over the 1950-2580 ms interval (cluster-based permutation test: *p* < 0.001; **Fig. 5E**), which falls within the 1-3 s post-stimulus period. Moreover, analysis of all contact pairs within this interval revealed that 38 out of 40 pairs exhibited face-specific modulation of *z*-scored dwPLI (face>shape; linear mixed-effects model: *t* = 4.32, *p* < 0.001; **Fig. 5F**). Given that the face-specific enhancement of *z*-scored dwPLI was primarily observed during the 2-3 sec interval, we further divided the 1-3 sec period into two sub-windows (1-2 sec and 2-3 sec) to examine whether directional information flow exhibited similar temporal specificity. Frequency-domain GC analysis was applied to both the l→m and m→l directions. Results showed that during 2-3 sec, there was a significant face-specific GC increase (face>shape) in the m→l direction within 2-16 Hz band (linear mixed-effects model: *t* = 2.09, *p* = 0.040; **Fig. 5G**). And this increase was significantly greater in the m→l direction compared to the reverse direction (linear mixed-effects model: *t* = 2.89, *p* = 0.005). No effects were observed during the 1-2 sec (all *ps* > 0.05). Taken together, these findings suggest that the medial amygdala is consistently engaged during face processing within 1-3 sec post-stimulus, and assumes a driving role in late-stage (>2 sec) information transfer.

### Intracranial stimulation of amygdala subregions differentially modulates fear and face processing

To further examine the causal role of amygdala subregions in fear and face processing, a new group of seven participants (see **Table 2**) received intracranial electrical stimulation (iES) of lAmyg (*N* = 4) or mAmyg (*N* = 5) sites during a separate EFMT task (**Fig. 6A**). In a random 50% of trials, 50-Hz stimulation trains (200 ms) were delivered time-locked to the image onset. The stimulation amplitude was determined individually based on the mapping procedure, with a mean current of 3.83 ± 1 mA across participants (**Fig. 6B**).To evaluate the site-specific behavioral effects of iES, separate linear mixed-effects models were fitted for the lAmyg and mAmyg stimulation sites. In each model, reaction time or accuracy served as the dependent variable, stimulation (stimulation *vs*. non-stimulation) and condition (fearful/neutral/shape) were included as fixed effects, and participant and trial number were modeled as random effects to account for repeated measurements and between-subject variability.

**Fig. 6.**
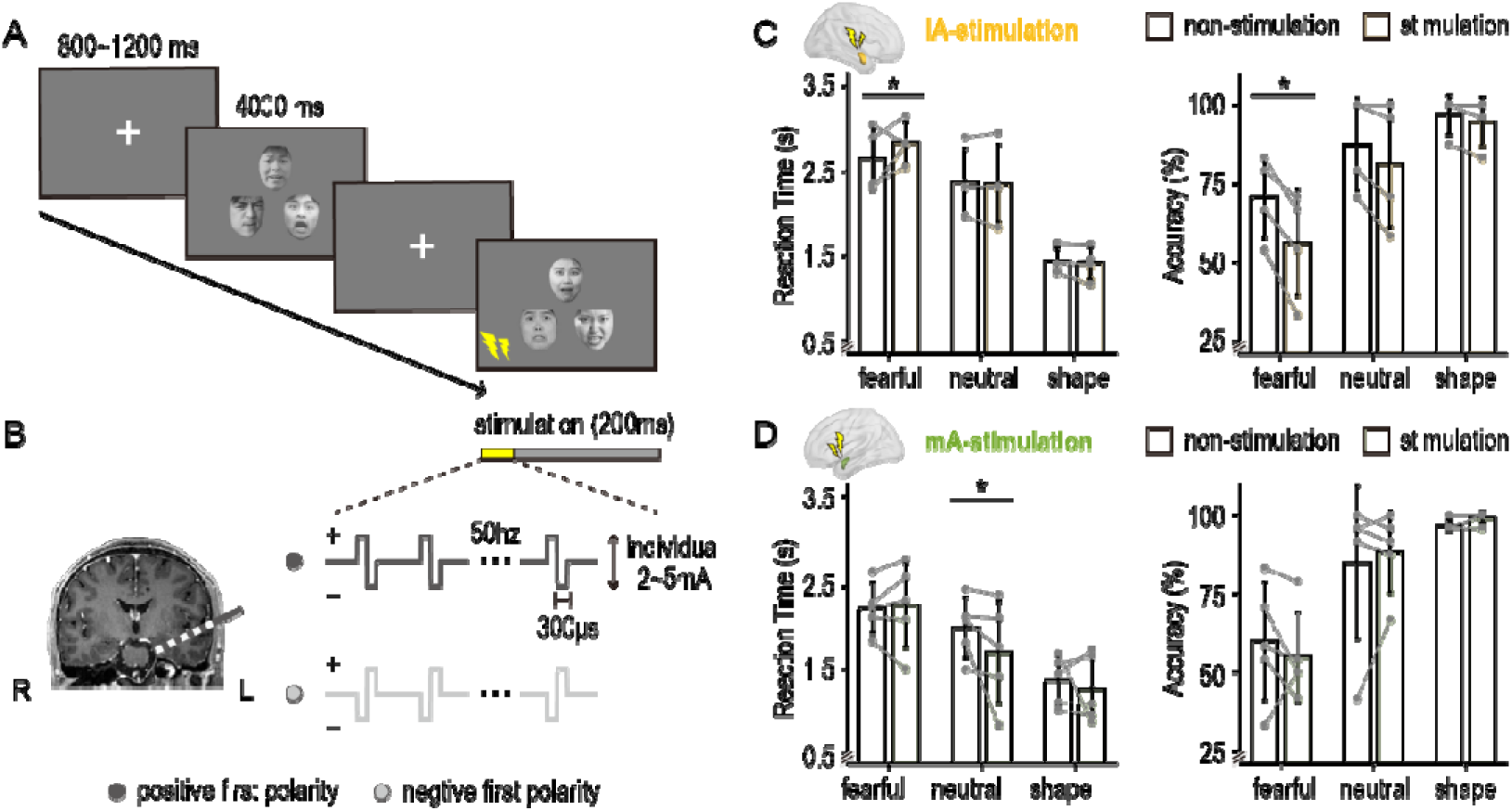
Intracranial stimulation of the lateral amygdala subregion disrupts behavioral performance during fear processing. (A) Example trials of the experimental task. In 50% of randomly selected trials, intracranial stimulation was delivered at image onset for 200 ms, with no more than three stimulation trials occurring consecutively. (B) Intracranial stimulation: biphasic currents delivered through adjacent contacts in the lateral or medial amygdala with 50-Hz stimulation trains (width 300μs). While the stimulation amplitude was determined individually based on the mapping procedure, with a 2-5mA range across participants. (C) Reaction time (left) and accuracy (right) during the lAmyg stimulation. The iES of the lAmyg selectively disrupted fear processing, marked by prolonged reaction times (linear mixed-effects model: *t* = 2.25, *p* = 0.025) and reduced recognition accuracy (*t* = 2.47, *p* = 0.014). Light yellow and yellow bars show means with standard deviation for no-stimulation and stimulation trials, separately. Gray dots indicate individuals and gray lines connect stimulation conditions. (D) Reaction time (left) and accuracy (right) during mAmyg stimulation. The iES to the mAmyg shortened reaction time to neutral faces (*t* = 1.99, *p* = 0.029). Symbols are consistent with panel C, with light green indicating no-stimulation trials and green indicating stimulation.

**Table 2.** Participants characteristics with intracranial electrical stimulation.

| Patient | Age | Sex | Stim region | Stim intensity |
| --- | --- | --- | --- | --- |
| 1 | 36 | male | lAmyg, right | 5mA |
| 2 | 34 | male | lAmyg, right | 4mA |
| 3 | 29 | male | lAmyg, right/mAmyg, right | 2mA/5mA |
| 4 | 34 | male | lAmyg, left/mAmyg, left | 4mA/4mA |
| 5 | 41 | male | mAmyg, right | 4mA |
| 6 | 15 | male | mAmyg, right | 4mA |
| 7 | 24 | male | mAmyg, right | 2.5mA |

Abbreviations: lAmyg, lateral amygdala; mAmyg, medial amygdala

For the lAmyg stimulation, we found that it caused a selective impairment in fear processing (**Fig. 6C**). Specifically, the iES prolonged mean reaction time of the fearful trials from 2.6 ± 0.7 sec (non-stimulation) to 2.8 ± 0.6 sec (stimulation; *t* = 2.25, *p* = 0.025), and reduced recognition accuracy from 69.8% ± 13.2% to 56.2% ± 16.8% (*t* = 2.47, *p* = 0.014). While it did not affect the other two conditions regardless of reaction time (neutral: *p* = 0.71; shape: *p* = 0.70) or accuracy (neutral: *p* = 0.22; shape: *p* = 0.68). By contrast, for the mAmyg stimulation, the effect of iES was primarily observed in reaction time to neutral faces. The iES significantly shortened the reaction time from 1.99 ± 0.36 sec (non-stimulation) to 1.83 ± 0.61 sec (stimulation; *t* = 1.99, *p* = 0.046; **Fig. 6D**). No effect was found in the fearful or shape conditions (fearful: *p* = 0.26; shape: *p* = 0.38). And no effect was found on recognition accuracy (all *ps* > 0.05). Thus, the iES perturbation provides causal evidence that the lateral and medial subregions have functional specialization in fear-specific and face-specific processing.

## Discussion

These convergent findings indicate that human amygdala subregions contribute to distinct, temporally dissociated components of fearful-expression recognition (**Fig. 7**). Our data suggest that the lateral subregion first differentiated fearful from neutral stimuli between ∼360 and 1110 ms post-stimulus, exhibiting higher decoding accuracy with enhanced theta/alpha-band power. This early response was followed by a unidirectional theta/alpha-band transmission from the lateral to the medial subregion, which showed delayed, sustained activation at a later stage (∼1750-3000 ms) with increased decoding accuracy and theta/alpha power. By contrast, during face encoding the medial subregion began to represent face-specific information in the 2-16 Hz at ∼1090 ms post-stimulus with superior decoding and spectral power, and subsequently conveyed this information back to the lateral subregion approximately 2 sec later, completing this process. Crucially, intracranial stimulation of either subregion produced selective modulation of behavioral responses to fear detection versus face recognition, supporting causal roles for these dynamics. Collectively, our study reveals that the lateral amygdala predominates in early fear detection while the medial amygdala drives later face encoding, and that their temporally structured interactions jointly shape fearful-expression recognition.

**Fig. 7.**
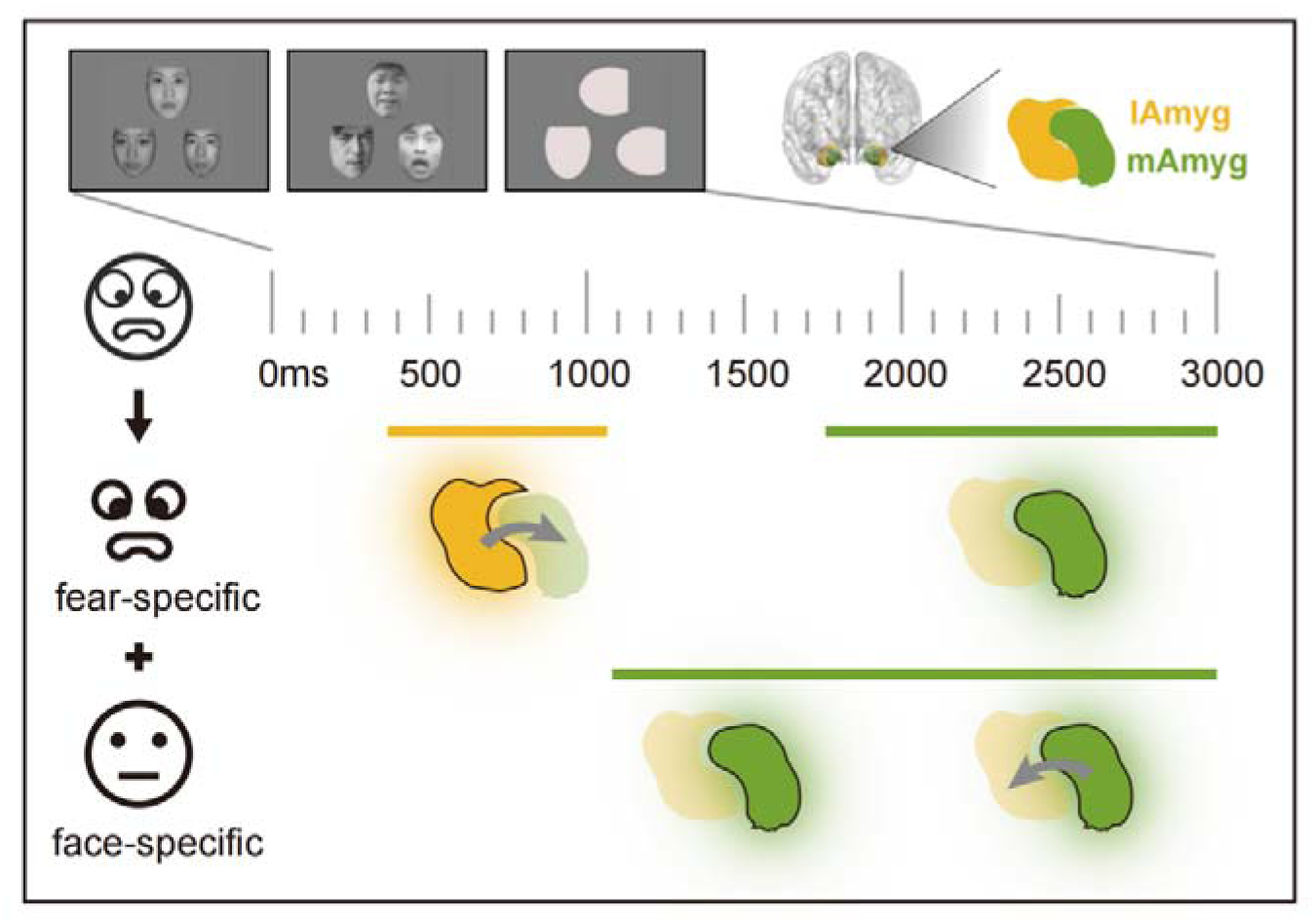
Summary and time course of neural activity of human amygdala subregions and information transmission during fearful-expression recognition. All data reported pertain to stimulus-locked responses during processing. Amygdala subregions involved in task-specific processing are indicated by black lines and glowing schematics. The temporal profiles of regional activation are represented by solid horizontal lines, indicating the onset and duration of observed processes in the lateral (lAmyg, yellow) and medial amygdala (mAmyg, green). Fearful-expresion recogntion not only depends on rapid fear detection, but also needs facial information encoding. Fearful-specific processing (top) was associated with early activation of theta/alpha-band (4-12 Hz) activity in the lAmyg (360-1110 ms), followed by enhanced phase synchronization and increased directional information flow from lAmyg to mAmyg, with solid arrows denoting significant directional influence. Then the mAmyg showed dominant sustained activity at later stage (1750-3000 ms). In contrast, face-specific processing (bottom) elicited delayed responses, with mAmyg activating from ∼1090 ms and relayed the low-frequency (2-16 Hz) activity back to the lAmyg after 2000 ms, which drives this process. The inset on the top illustrates the task context corresponding to the data analysis and the anatomical locations of lAmyg and mAmyg.

In rodents, the BLA receives rapid thalamic and cortical sensory input and forms the primary representation of threat cues ^19,21^, whereas the CeA occupy an output position that drives brainstem and hypothalamic defensive programs ^33,34^. Cross-species work documents subregional functional specialisation in fear processing ^11,18^, supporting a conserved division of labour. Human fMRI comparisons of amygdala subregions have produced mixed results, partly because BOLD is slow and indirect, reporting variable medial versus lateral dominance ^23,24,26^. Whereas an iEEG study revealed rapid (∼200 ms), stimulus-locked amygdala responses ^4^ but typically treat the structure as unitary. Extending these lines, our study highlights the temporal dynamics of subregional engagement, in which the lateral subregion predominates in early threat detection, whereas the medial subregion shows delayed and sustained activation at later stage. Critically, intracranial stimulation of the lateral subregion impaired behavioral performance on fear-detection, providing causal evidence for this functional dissociation. This result aligns with prior reports that amygdala stimulation can disrupt emotional memory encoding ^35^.

Further, directed analyses revealed an early, preferential routing of threat-related signals from the lateral to the medial amygdala, carried predominantly in the theta/alpha band. This intra-amygdalar feedforward pathway is anatomically plausible given direct BLA→CeA projections that support rapid fear responses in animals ^18,21^, and it is indirectly supported in humans by lesion data showing disrupted CeA-brainstem coordination after BLA damage ^19^. Although prior work has largely emphasized amygdala interactions with distributed cortical and subcortical targets ^36,37^, our study specifically characterizes frequency-specific, within-amygdala routing and shows that this pathway dynamically coordinates early threat detection. Additionally, the dominance of theta/alpha rhythms matches literature implicating low-frequency oscillations in amygdala-prefrontal coupling during fear encoding ^38^ and in amygdala-hippocampal coordination for negative-affect processing ^39^. We extend this framework by demonstrating that theta/alpha activity serves not only as a carrier for long-range communication but also as a conserved mechanism for intra-amygdalar signalling. Collectively, time-resolved electrophysiology and causal perturbation indicate that temporal dissociation and frequency-specific interactions within the amygdala provide a parsimonious mechanism for rapid fearful-expression recognition.

Unlike rapid fear detection, face processing in our data is dominated by the medial amygdala. Primate studies report that amygdala neurons encode non-emotional facial dimensions ^40^ and that these representations are subregionally organised, with the basolateral complex shows strong visual-feature selectivity and centromedial groups are more closely involved in gaze, attention and autonomic output ^41–43^. Human single-unit studies similarly implicate mid-medial sites in encoding familiarity, identity and other high-level facial cues ^7^. These convergent findings support our observation that the medial subregion predominates in face processing. Moreover, our data show that this medial dominance is dynamic, with the medial subregion beginning to encode face information after 1 sec post-stimulus and sustaining low-frequency activity throughout the process.

While a prior iEEG study reported an early gaze-sensitive N200 activity to neutral face stimuli in anterior-medial amygdala electrodes ^44^, the later signal we observe may reflect attention-gated, higher-order feature analysis rather than automatic face detection ^42,45^. First, the dense connections of the medial/centromedial nuclei with subcortical and paralimbic targets (e.g., brainstem, insula) position them to modulate arousal and attentional state via autonomic pathways ^10,33,46^. Accordingly, the medial→ lateral information flow we observe at approximately 2 sec may reflect a feedback process in which the medial subregion conveys attention-enhanced facial representations to the lateral subregion, facilitating detailed feature analysis and completion of face encoding. Second, face-specific responses occupy a broad 2-16 Hz range, a spectral profile that could support multi-frequency integration. Lower-theta components (∼2-6 Hz) may bind temporal feature elements, whereas higher-alpha components (∼8-16 Hz) may subserve attentional selection ^29^. Thus, these evidence suggest that the medial amygdala driving late-stage face encoding may operate through an attention-gated, integrative mechanism.

One further issue merits discussion. Identical intracranial electrical stimulation produced opposite behavioral effects when applied to medial versus lateral amygdala, with lateral stimulation impairing fear detection while medial stimulation facilitated face recognition. Several possible explanations could account for this divergence. First, medial sites lie close to output pathways and can therefore more directly influence arousal and behavioral readiness ^33^, which is consistent with reports that brief amygdala stimulation can enhance encoding of neutral images without eliciting subjective emotion ^47^. Second, the behavioral effect of stimulation may depend on its timing relative to the region’s endogenous recruitment, because pulses delivered before medial engagement could boost excitability and thereby amplify subsequent information integration in a state-dependent manner ^48,49^. Third, intracranial stimulation can produce facilitative or disruptive network effects, which depend on stimulation site, timing and the ongoing network state ^50^. Fine spatial heterogeneity and current spread may also produce different outcomes even between nearby contacts ^51^. Therefore, the contrasting stimulation effects provide causal evidence that the medial amygdala contributes to late-stage face processing and that its temporal recruitment is behaviorally meaningful. Future studies using larger cohorts and systematic variation of stimulation timing and parameters are needed to disentangle these spatial and temporal causal dynamics.

In sum, our data provide converging neural and behavioral evidence for a temporally structured, bidirectional interaction between human amygdala subregions to support distinct cognitive components of fearful-expression recognition. The lateral subregion predominates in early fear detection, whereas the medial subregion drives later face-feature integration. While the amygdala has long been implicated in fear processing, we link human circuit dynamics to animal models of amygdaloid subdivision, thereby broadening mechanistic accounts of how the amygdala implements multidimensional processing in humans.

## Methods

### Participants

Twelve drug-resistant epilepsy patients (3 females; mean age ± SEM: 32±12 years) participated in this study. All of them were undergoing iEEG monitoring at the Department of Neurosurgery, Chinese PLA General Hospital to localize epileptic foci for potential surgical resection. Stereotactic depth EEG electrodes (0.8 mm diameter, 2 mm length, 1.5 mm spacing; Sinovation Medical Technology Co., Ltd., Beijing, China) were stereotactically implanted. Electrode placement was determined solely based on clinical requirements. We included all patients that accepted to participate in the study and that had electrodes implanted in the amygdala. We listed the patients’ demographic information, number of contacts in the amygdala subregions and seizure onset zone in **Table 1**. Prior to testing, all participants provided written informed consent for the study, and the experimental procedures were approved by the ethics committee of the Chinese PLA General Hospital (No. S2021-394-02). There were no seizures recorded during any of the epochs, and any epochs with interictal epileptiform activity were excluded from analysis.

### Task

We employed an emotional face-matching task adapted from the block-design paradigm by Hariri et al ^52^. Participants completed three blocks in a fixed sequence: fearful-face, neutral-face, and shape-matching. Each block included 24 trials, with each trial beginning with a fixation cross (0.8-1.2 sec), followed by three images presented for 4 sec. Participants were instructed to select one of the two bottom images that matched the top image based on emotional expression (fearful-face condition), gender (neutral-face condition), or orientation (control-shape condition). All face images were drawn from the Chinese Facial Affective Picture System ^53^. The task was programmed and presented using MATLAB (The MathWorks) with Psychtoolbox-3 (PTB-3; Brainard & Pelli) .

### Depth electrode localization

Electrode contact localization was conducted using post-implantation computed tomography (CT) scans and structural T1-weighted magnetic resonance imaging (MRI). CT scans were co-registered to the post-implantation MRI for each patient using Brainstorm software ^54^, and electrode contacts were visually identified on the co-registered CT-MRI images. Contact positions were further verified by the neurosurgeon (Y.Z.) through inspection of merged pre-operative MRI and post-implantation CT images along the electrode trajectory. Contact coordinates in native space were transformed into MNI space. Amygdala contacts were identified based on the Brainnetome Atlas ^55^ and classified into medial and lateral subregions. The final dataset included 58 amygdala contacts, comprising 38 contacts in the lateral subregion and 20 contacts in the medial subregion across all participants. On average, there were 3.2 D±D 1.9 lateral contacts per participant (range: 1-6; 12 of 12 participants) and 1.7D±D1.6 medial contacts per participant (range: 0-4; 7 of 12 participants). Data from 7 participants who had contacts in both subregions were included in the inter-regional connectivity analyses. The seizure onset zones (SOZs) of all patients are summarized in **Table 1**, with no SOZ involving the amygdala. **Fig. 2D** shows electrode locations transformed into standard MNI space, overlaid on the amygdala subregions across all participants.

### Data acquisition and preprocessing

Intracranial EEG data were recorded using the NeuSen H high-density neural signal acquisition system (Neuracle Technology Co., Ltd., Beijing, China) with a band-pass filter of 0.03–500 Hz and a sampling rate of 2000 Hz. The recordings were subsequently downsampled to 1 kHz and band-pass filtered between 1 and 200 Hz using a zero-phase finite impulse response filter with a Hamming window. Line noise (50 Hz) and its harmonics were removed via discrete Fourier transform. The preprocessed data were visually inspected to identify and exclude channels containing epileptiform activity or artifacts. The remaining signals were then re-referenced to a white matter contact located distant from the amygdala. Data were then segmented into event-related epochs of 4500 ms, which were time-locked to stimulus onset and spanned -500 to 4,000 ms. We rejected epochs with residual artifacts by visual inspection (107/864 trials, 12.38%). We performed preprocessing routines with FieldTrip ^56^, EEGLAB ^57^, and custom scripts in MATLAB R2018b (the MathWorks, Natick, MA, USA).

### Time-frequency analysis

Time-frequency power was computed for each contact within the medial and lateral subregions at each trial in each patient. Broadband signals were convolved with complex-valued Morlet wavelets (6 cycles) to extract power information from 1 to 150 Hz in 1 Hz increments with a time resolution of 1 ms. Task-related power was normalized on a trial-by-trial basis using a statistical bootstrapping approach, as established in previous studies ^58^. Briefly, for each contact and frequency, a null distribution was generated by randomly sampling and averaging data points from the 500 ms pre-stimulus baseline, with this process repeated 1,000 times. Raw power at each time point during the task was then *z*-scored by comparing it to this null distribution to generate the *z*-scored power for all conditions. The task-induced power was considered significant when the *z*-scored power exceeded the value of 1.96 (*p* < 0.05).

### Responsive contacts selection

Responsive contacts were defined as those exhibiting *z*-scored power values exceeding 1.96 for at least 50 ms, as reported in a previous study ^29^. Contacts were identified separately for each condition (fearful, neutral, shape) and frequency band (theta [4-7 Hz], alpha [8-12 Hz], beta [13-30 Hz], gamma [31-60 Hz] and high-gamma [61-150 Hz]). For both subregions, the fearful condition elicited more responsive contacts than the other two conditions across all bands (**Fig. S1B**).

Additionally, responsive contacts under the neutral and shape conditions largely overlapped with those identified in the fearful condition. Given this convergence, we defined the union of contacts responsive in any condition across frequency bands as the set for subsequent analyses. This yielded 17 responsive contacts in the mAmyg and 31 contacts in the lAmyg.

### Time-resolved decoding analysis

We performed a time-resolved multivariate decoding analysis to examine how amygdala subregions distinguish fear-specific (fearful vs. neutral) or face-specific (neutral vs. shape) processing as a function of time. The support vector machine (SVM) is widely utilized in decoding analysis in neuroimaging studies ^59,60^, particularly due to its suitability for datasets with relatively small sample sizes. Hence, our analysis was performed based on a linear SVM as the classifier via the LIBSVM package ^61^ in MATLAB. We conducted binary classification using *z*-scored time-frequency power features. To increase the impact of decoding analysis on a larger population, for each condition, the *z*-scored power at the trial level for all participants were merged as the data (fearful: 250 samples, neutral: 247 samples, shape: 260 samples) used in the classification.

Taking the decoding of fearful-faces versus neutral-faces as an example, for each subregion, the theta/alpha-band (4-12 Hz) power features were extracted for each trial and participant. The analysis window (0-3000 ms post-stimulus) was divided into overlapping 100 ms time windows with a 10 ms step size. Within each window, the *z*-scored power features for each trial included 9×100 = 900 values (where 9 denotes the frequency range from 4 to 12 Hz and 100 denotes the time points), which were then converted into a one-dimensional feature vector. Then, we split 70% of the data from each condition, merging them across both conditions to form the training dataset, while the remaining data served as the testing dataset. To reduce feature dimensionality, principal component analysis (PCA) was applied to the training dataset, retaining several principal components (K components: 10.1 ± 0.9, ranging from 7 to 12) that explained at least 99% of the variance. The testing dataset was then transformed using the PCA matrix fitted to the training data. We trained the SVM classifier with a linear kernel with a cost equal to one. This process was repeated 100 times for cross-validation, as in previous studies ^60,62^. The accuracy of the classifier as a performance measure was averaged across 100 cross-validations. This procedure was repeated across all time windows (311 windows). The schematic of the time-resolved decoding analysis steps is shown in **Fig. 3B**. In addition, we also decoded neutral-faces versus control-shapes using *z*-scored power features from both subregions. Aside from the differences in sample size and the power features from 2-16 Hz frequency range, all other steps were the same as described above.

### Debiased weighted phase lag index analysis

To examine whether potential interactions occurs between the amygdala subregions, phase synchronization was quantified by computing the dwPLI across all contact pairs spanning the two subregions. The dwPLI quantifies non-zero phase lags through the asymmetry of phase-difference distributions and reduces bias from volume conduction, noise, and limited sample size ^32^. The dwPLI takes values between 0 and 1, with values approaching 1 reflecting stronger phase synchrony between signals. We computed the dwPLI in the time-frequency domain from 1-150 Hz in 1 Hz step for each contact pair within the same hemisphere (one contact from the medial and the other from the lateral subregion) for the trials under each condition. Each signal was convolved with a family of complex Morlet wavelets (wavelet width logarithmically increasing from 4 to 8 cycles) to obtain the analytic representation 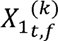 and 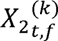 at trial *k*. At each time-frequency point (*t*, *f*), we extracted the imaginary part of the cross-spectrum:

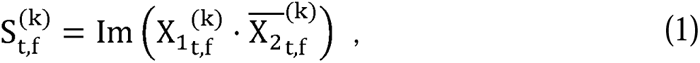

and estimated dwPLI given by ^63^:

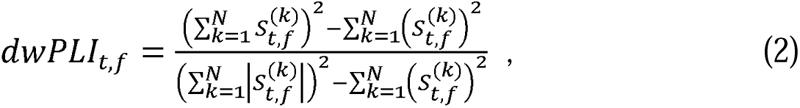

where *N* denotes the number of trials.

To evaluate the statistical significance of dwPLI, a null distribution was created by randomly shuffling the trials for each contact pair and computing the corresponding dwPLI spectrogram and repeating this procedure for 100 times. The original dwPLI values were then *z*-scored against the null distribution computed separately for each condition. Significant dwPLI was thresholded at |*z*| > 1.96 (*p* < 0.05).

### Granger causality analysis

After establishing the phase synchrony, which measures undirected connectivity between the amygdala subregions, we proceeded to investigate the directionality of their interaction using nonparametric frequency-domain Granger causality. This approach measures the degree to which the signal from a region (i.e., the lAmyg) can be better predicted by incorporating information from another signal (i.e., the mAmyg) in a specific frequency band, and vice versa, which is suitable for short time windows and non-stationary data ^17,64^. For each contact pair between the two subregions, the signals from the condition-specific time windows of interest were extracted (for fear-specific analysis, early (360-1110 ms) and late (1750-3000 ms) windows were selected; for face-specific analysis, the 1-3 sec were selected).

We rearranged the multichannel iEEG data into a three-dimensional matrix (time × trial × channel), directly applied the multitaper Fourier method (2 × NW tapers, NW = 2) to estimate power and cross-spectra, and obtained the spectral density matrix by averaging across trials. The transfer function *H*(*f*) and the noise covariance Σ were then derived through Wilson spectral factorization, such that

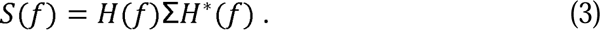

In the frequency domain, we computed the GC index according to the following formulation ^65^:

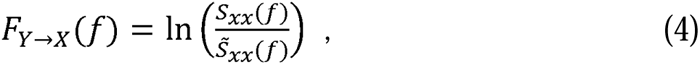

here *S_xx_*(*f*) denotes the total power spectrum of the effect variable *X*, and *S^∼^_xx_*(*f*) represents the intrinsic power obtained by setting the contribution from *Y* to zero and reconstructing it through the transfer function *H*(*f*). When *F_Y_*→*_X_*(*f*) > 0, it indicates the presence of directional information flow from Y to X within frequency band *f*.

We computed directional causality spectra separately for medial→lateral and lateral → medial amygdala, and averaged across contact pairs to estimate the overall strength of information flow between subregions. To assess the statistical significance of the GC index, we generated a null distribution by randomly shuffling the signals between contact pairs 100 times and then averaging across all pairs. Original values exceeding the 95th percentile of this null distribution were considered significant and retained for further analyses. Finally, based on the observations from the dwPLI analyses, we extracted the average GC index within specific frequency bands of interest (4-12 Hz for fear-specific processing; 2-16 Hz for face-specific processing).

### Intracranial stimulation during emotional face-matching task

As the iEEG task assessed the involvement of amygdala subregions in fear and face processing, we next aimed to examine the causal contributions of the lateral and medial subregions via direct electrical stimulation. To do so, we recruited 7 additional participants (see **Table 2**) to perform an emotional face-matching task while receiving the iES to a targeted pair of subregion electrode contacts. Before the task, the safe stimulation amplitudes were determined by a neurologist (Y.Z.) using an iterative mapping procedure. Stimulation was applied only after a neurologist identified safe amplitudes using an iterative mapping procedure. Mapping was performed in 0.5 mA increments and monitored for after-discharges. The final stimulation intensity was set to 80% of the after-discharge threshold or capped at 5 mA, ensuring that no discomfort or seizures were induced. Individual stimulation intensities are listed in **Table 2**.

Then, the iES was delivered in a bipolar configuration through a single pair of adjacent contacts. Each stimulation consisted of a 200-ms train of charge-balanced biphasic rectangular pulses (300 µs pulse width) applied continuously at 50 Hz frequency (**Fig. 6B**). Stimulation was time-locked to image onset while participants responded to fearful-face, neutral-face, or control-shape conditions (**Fig. 6A**). Each condition comprised 48 trials, with stimulation applied to 50% of trials in a pseudo-random order (no more than three consecutive stimulated trials). Of the seven participants, 4 received the iES in the lAmyg and 5 in the mAmyg. Among these participants, two underwent the iES in both subregions. The two stimulation procedures were conducted on separate days, separated by at least a 24-hour interval.

### Statistical analysis

For time-frequency analyses, we compared time-frequency *z*-scored power between fearful and neutral trials at the contact level using a cluster-based permutation test ^66^ to identify fear-specific effect within each subregion; an analogous procedure (neutral-face vs. control-shape) was applied to evaluate face-specific effect. In addition, we extracted mean *z*-scored power within predefined time windows and frequency band for each subregional contact and condition, and tested the differences between conditions / subregions using linear mixed-effect (LME) models at contact level. Specifically, in fear-specific analyses, the condition effect were tested with a model including *Condition* (fearful vs. neutral) as the fixed effect, with *Power* as the dependent variable, and *Subject* and *Contact* as random effects, the model was structured as follows:

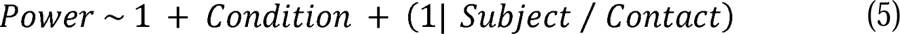

The Regional effect was assessed with a model including *Region* (lAmyg vs. mAmyg) and *Time* (early vs. late) as fixed effects, with same dependent variable and random-effects:

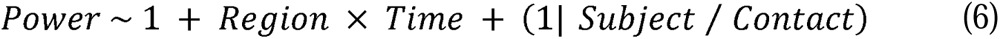

In the face-specific analyses, only one time window was considered. In contrast, the regional model included Region as the fixed effect, with the dependent variable and random-effects structure unchanged. The conditional model adopted the same structure as the fear-specific analyses.

For time-resolved decoding analyses, we used a non-parametric permutation test to examine the significance of decoding accuracy. At each time window, we randomly shuffled the relationship between labels and data 100 times to generate a null distribution of decoding accuracy, as consistent with our previous study ^60^. The true decoding accuracy was then compared to this null distribution, with values exceeding the 95^th^ percentile (*p* < 0.05) considered significant. For subregional comparisons, we adopted paired t-tests on decoding accuracy from the two subregions with 100 cross-validation iterations as samples.

To identify when the time-varying *z*-scored dwPLI showed condition-specific differences, we extracted the frequency-specific values and compared them between conditions using a cluster-based permutation test across time. Within the time window containing a significant cluster, we further quantified the effect at the contact-pair level by computing the contrast between conditions for each pair and testing these contrasts against zero with a LME model including *Condition* as the fixed effect, with fear-specific (or face-specific) *dwPLI* as the dependent variable, and *Subject* and *Contact Pair* as random effects. In addition, for the statistical analyses of the GC index, we extracted the conditional contrasts at the contact-pair level within predefined time windows. These contrasts were then used as the dependent variables in LME models, which followed similar structural specification as Equations (5) and (6) above, to assess conditional and directional effects.

To analyze the behavioral effects induced by iES, trial-level data were analyzed separately for tasks involving lAmyg or mAmyg stimulation, using linear mixed-effects models. We tested reaction time and recognition accuracy as separate dependent variables. For each model, Stimulation (stimulation/non-stimulation) and Condition (fearful/neutral/shape) served as fixed effects, with either reaction time or accuracy as the dependent variable. Subjects and trial number served as random effects.

## Data availability

The preprocessed data generated in this study have been deposited in the Open Science Framework (OSF) database at https://osf.io/apxj4/, and will be available upon publication.

## Code Availability

The custom code supporting this study is available at https://github.com/lxdev-bit/human-amygdala-subregions-iEEG. Standard preprocessing of iEEG data was performed using EEGLAB and FieldTrip.

## Acknowledgement

This work received support from the following sources: Beijing Natural Science Foundation (No.5244049 to D.C.; No.L256006 to J.L.), National Natural Science Foundation of China (No. 32400883 to D.C.; No. 32271085 to J.L.; No. 82001798 to Y.Z.), the Young Talent Project of Chinese PLA General Hospital (No. 20230403 to Y.Z.), the Innovation Incubation Project of Chinese PLA General Hospital for the Department of Neurosurgery (No. SWXB-FH-004 to Y.Z.) and Open Research Fund of the State Key Laboratory of Cognitive Neuroscience and Learning (CNLYB2301 to D.C.).

## Author contributions

Conceptualization, Y.Z., J.Z. and X.Y.; Methodology, D.C. and J.C.; Data Collection, Y.Z., Y.D., and H.Z.; Formal Analysis: D.C., and J.C.; Writing- Original Draft, D.C., and J.C.; Writing-Review & Editing, D.C., J.C., J.L., and Y.Z.; Funding Acquisition, D.C., Y.Z. J.L., and J.Z.; Resources, Y.Z. and J.Z.; Supervision, Y.Z. and D.C.

## Declaration of Large Language Models in the writing process

This manuscript was prepared with the assistance of ChatGPT to improve linguistic readability. The authors thoroughly reviewed and revised all content as necessary.

## Reference

1 Adolphs, R. Fear, faces, and the human amygdala. Curr Opin Neurobiol 18, 166–172 (2008). 10.1016/j.conb.2008.06.006

2 Sah, P. Fear, anxiety, and the amygdala. Neuron 96, 1–2 (2017).

3 LeDoux, J. Rethinking the emotional brain. Neuron 73, 653–676 (2012).

4 Mendez-Bertolo, C. et al. A fast pathway for fear in human amygdala. Nat Neurosci 19, 1041–1049 (2016). 10.1038/nn.4324

5 Wang, Y. et al. Rapid Processing of Invisible Fearful Faces in the Human Amygdala. J Neurosci 43, 1405–1413 (2023). 10.1523/JNEUROSCI.1294-22.2022

6 Cao, R. et al. A neuronal social trait space for first impressions in the human amygdala and hippocampus. Molecular Psychiatry 27, 3501–3509 (2022).

7 Cao, R. et al. Feature-based encoding of face identity by single neurons in the human amygdala and hippocampus. Nature Human Behaviour, 1–16 (2025).

8 Rutishauser, U., Mamelak, A. N. & Adolphs, R. The primate amygdala in social perception - insights from electrophysiological recordings and stimulation. Trends Neurosci 38, 295–306 (2015). 10.1016/j.tins.2015.03.001

9 Gothard, K. M. Multidimensional processing in the amygdala. Nat Rev Neurosci 21, 565–575 (2020). 10.1038/s41583-020-0350-y

10 Bzdok, D., Laird, A. R., Zilles, K., Fox, P. T. & Eickhoff, S. B. An investigation of the structural, connectional, and functional subspecialization in the human amygdala. Human brain mapping 34, 3247–3266 (2013).

11 Hochgerner, H. et al. Neuronal types in the mouse amygdala and their transcriptional response to fear conditioning. Nat Neurosci 26, 2237–2249 (2023). 10.1038/s41593-023-01469-3

12 Pessoa, L., Medina, L., Hof, P. R. & Desfilis, E. Neural architecture of the vertebrate brain: implications for the interaction between emotion and cognition. Neuroscience & Biobehavioral Reviews 107, 296–312 (2019).

13 Saygin, Z. M., Osher, D. E., Augustinack, J., Fischl, B. & Gabrieli, J. D. Connectivity-based segmentation of human amygdala nuclei using probabilistic tractography. Neuroimage 56, 1353–1361 (2011).

14 Avecillas-Chasin, J. M. et al. Connectivity-based parcellation of the amygdala and identification of its main white matter connections. Scientific reports 13, 1305 (2023).

15 Solano-Castiella, E. et al. Diffusion tensor imaging segments the human amygdala in vivo. Neuroimage 49, 2958–2965 (2010).

16 Sylvester, C. M. et al. Individual-specific functional connectivity of the amygdala: A substrate for precision psychiatry. Proceedings of the National Academy of Sciences 117, 3808–3818 (2020).

17 Sawada, M. et al. Mapping effective connectivity of human amygdala subdivisions with intracranial stimulation. Nat Commun 13, 4909 (2022). 10.1038/s41467-022-32644-y

18 Kim, J., Zhang, X., Muralidhar, S., LeBlanc, S. A. & Tonegawa, S. Basolateral to central amygdala neural circuits for appetitive behaviors. Neuron 93, 1464–1479. e1465 (2017).

19 Terburg, D. et al. The Basolateral Amygdala Is Essential for Rapid Escape: A Human and Rodent Study. Cell 175, 723–735 e716 (2018). 10.1016/j.cell.2018.09.028

20 Adolphs, R. et al. A mechanism for impaired fear recognition after amygdala damage. Nature 433, 68–72 (2005).

21 Liu, J., Totty, M. S., Bayer, H. & Maren, S. Integrating Aversive Memories in the Basolateral Amygdala. Biol Psychiatry (2025). 10.1016/j.biopsych.2025.03.019

22 Webler, R. D. et al. The neurobiology of human fear generalization: meta-analysis and working neural model. Neuroscience & Biobehavioral Reviews 128, 421–436 (2021).

23 Hrybouski, S. et al. Amygdala subnuclei response and connectivity during emotional processing. Neuroimage 133, 98–110 (2016).

24 Labuschagne, I. et al. Specialization of amygdala subregions in emotion processing. Hum Brain Mapp 45, e26673 (2024). 10.1002/hbm.26673

25 Boll, S., Gamer, M., Kalisch, R. & Büchel, C. Processing of facial expressions and their significance for the observer in subregions of the human amygdala. Neuroimage 56, 299–306 (2011).

26 Hurlemann, R. et al. Segregating intra-amygdalar responses to dynamic facial emotion with cytoarchitectonic maximum probability maps. Journal of neuroscience methods 172, 13–20 (2008).

27 Hariri, A. R., Tessitore, A., Mattay, V. S., Fera, F. & Weinberger, D. R. The amygdala response to emotional stimuli: a comparison of faces and scenes. Neuroimage 17, 317–323 (2002). 10.1006/nimg.2002.1179

28 Wright, P. & Liu, Y. Neutral faces activate the amygdala during identity matching. Neuroimage 29, 628–636 (2006). 10.1016/j.neuroimage.2005.07.047

29 Helfrich, R. F. et al. Neural mechanisms of sustained attention are rhythmic. Neuron 99, 854–865. e855 (2018).

30 Xia, M., Wang, J. & He, Y. BrainNet Viewer: a network visualization tool for human brain connectomics. PloS one 8, e68910 (2013).

31 Grootswagers, T., Wardle, S. G. & Carlson, T. A. Decoding dynamic brain patterns from evoked responses: a tutorial on multivariate pattern analysis applied to time series neuroimaging data. Journal of cognitive neuroscience 29, 677–697 (2017).

32 Vinck, M., Oostenveld, R., van Wingerden, M., Battaglia, F. & Pennartz, C. M. An improved index of phase-synchronization for electrophysiological data in the presence of volume-conduction, noise and sample-size bias. Neuroimage 55, 1548–1565 (2011). 10.1016/j.neuroimage.2011.01.055

33 Janak, P. H. & Tye, K. M. From circuits to behaviour in the amygdala. Nature 517, 284–292 (2015).

34 Tye, K. M. et al. Amygdala circuitry mediating reversible and bidirectional control of anxiety. Nature 471, 358–362 (2011).

35 Qasim, S. E., Mohan, U. R., Stein, J. M. & Jacobs, J. Neuronal activity in the human amygdala and hippocampus enhances emotional memory encoding. Nat Hum Behav 7, 754–764 (2023). 10.1038/s41562-022-01502-8

36 Dal Monte, O., Chu, C. C., Fagan, N. A. & Chang, S. W. Specialized medial prefrontal–amygdala coordination in other-regarding decision preference. Nature neuroscience 23, 565–574 (2020).

37 Klein-Flügge, M. C. et al. Relationship between nuclei-specific amygdala connectivity and mental health dimensions in humans. Nature human behaviour 6, 1705–1722 (2022).

38 Chen, S. et al. Theta oscillations synchronize human medial prefrontal cortex and amygdala during fear learning. Science advances 7, eabf4198 (2021).

39 Zheng, J. et al. Amygdala-hippocampal dynamics during salient information processing. Nature communications 8, 14413 (2017).

40 Gothard, K. M., Battaglia, F. P., Erickson, C. A., Spitler, K. M. & Amaral, D. G. Neural responses to facial expression and face identity in the monkey amygdala. J Neurophysiol 97, 1671–1683 (2007). 10.1152/jn.00714.2006

41 Morrow, J., Mosher, C. & Gothard, K. Multisensory neurons in the primate amygdala. Journal of Neuroscience 39, 3663–3675 (2019).

42 Mosher, C. P., Zimmerman, P. E. & Gothard, K. M. Response characteristics of basolateral and centromedial neurons in the primate amygdala. J Neurosci 30, 16197–16207 (2010). 10.1523/JNEUROSCI.3225-10.2010

43 Mosher, C. P., Zimmerman, P. E. & Gothard, K. M. Neurons in the monkey amygdala detect eye contact during naturalistic social interactions. Current Biology 24, 2459–2464 (2014).

44 Huijgen, J. et al. Amygdala processing of social cues from faces: an intracrebral EEG study. Soc Cogn Affect Neurosci 10, 1568–1576 (2015). 10.1093/scan/nsv048

45 Minxha, J. et al. Fixations gate species-specific responses to free viewing of faces in the human and macaque amygdala. Cell reports 18, 878–891 (2017).

46 Pessoa, L. & Adolphs, R. Emotion processing and the amygdala: from a’low road’to’many roads’ of evaluating biological significance. Nature reviews neuroscience 11, 773–782 (2010).

47 Inman, C. S. et al. Direct electrical stimulation of the amygdala enhances declarative memory in humans. Proceedings of the National Academy of Sciences 115, 98–103 (2018).

48 Ezzyat, Y. et al. Direct brain stimulation modulates encoding states and memory performance in humans. Current biology 27, 1251–1258 (2017).

49 Merkow, M. B. et al. Stimulation of the human medial temporal lobe between learning and recall selectively enhances forgetting. Brain Stimulation 10, 645–650 (2017).

50 Mohan, U. R. & Jacobs, J. Why does invasive brain stimulation sometimes improve memory and sometimes impair it? PLoS Biol 22, e3002894 (2024). 10.1371/journal.pbio.3002894

51 Jacobs, J., Lega, B. & Anderson, C. Explaining how brain stimulation can evoke memories. Journal of Cognitive Neuroscience 24, 553–563 (2012).

52 Hariri, A. R., Mattay, V. S., Tessitore, A., Fera, F. & Weinberger, D. R. Neocortical modulation of the amygdala response to fearful stimuli. Biol Psychiatry 53, 494–501 (2003). 10.1016/s0006-3223(02)01786-9

53 Gong, X., Huang, Y.-X., Wang, Y. & Luo, Y.-j. Revision of the Chinese facial affective picture system. Chinese mental health journal (2011).

54 Tadel, F., Baillet, S., Mosher, J. C., Pantazis, D. & Leahy, R. M. Brainstorm: A user-friendly application for MEG/EEG analysis. Computational intelligence and neuroscience 2011, 879716 (2011).

55 Fan, L. et al. The human brainnetome atlas: a new brain atlas based on connectional architecture. Cerebral cortex 26, 3508–3526 (2016).

56 Oostenveld, R., Fries, P., Maris, E. & Schoffelen, J.-M. FieldTrip: open source software for advanced analysis of MEG, EEG, and invasive electrophysiological data. Computational intelligence and neuroscience 2011, 156869 (2011).

57 Delorme, A. & Makeig, S. EEGLAB: an open source toolbox for analysis of single-trial EEG dynamics including independent component analysis. Journal of neuroscience methods 134, 9–21 (2004).

58 Li, J. et al. Anterior–posterior hippocampal dynamics support working memory processing. Journal of Neuroscience 42, 443–453 (2022).

59 Tanigawa, H. et al. Decoding distributed oscillatory signals driven by memory and perception in the prefrontal cortex. Cell reports 39 (2022).

60 Yang, J. et al. Enhanced role of the entorhinal cortex in adapting to increased working memory load. Nat Commun 16, 5798 (2025). 10.1038/s41467-025-60681-w

61 Chang, C.-C. & Lin, C.-J. LIBSVM: A library for support vector machines. ACM transactions on intelligent systems and technology (TIST) 2, 1–27 (2011).

62 Li, J. et al. Functional specialization and interaction in the amygdala-hippocampus circuit during working memory processing. Nature Communications 14, 2921 (2023).

63 Cohen, M. X. Analyzing neural time series data: theory and practice. (MIT press, 2014).

64 Pagnotta, M. F., Dhamala, M. & Plomp, G. Benchmarking nonparametric Granger causality: Robustness against downsampling and influence of spectral decomposition parameters. NeuroImage 183, 478–494 (2018).

65 Dhamala, M., Rangarajan, G. & Ding, M. Analyzing information flow in brain networks with nonparametric Granger causality. Neuroimage 41, 354–362 (2008). 10.1016/j.neuroimage.2008.02.020

66 Maris, E. & Oostenveld, R. Nonparametric statistical testing of EEG-and MEG-data. Journal of neuroscience methods 164, 177–190 (2007).

