## Supplemental figure 1 and 2 for "Functional specialization and dynamical interaction in human amygdala subregions support fearful-expression recognition"

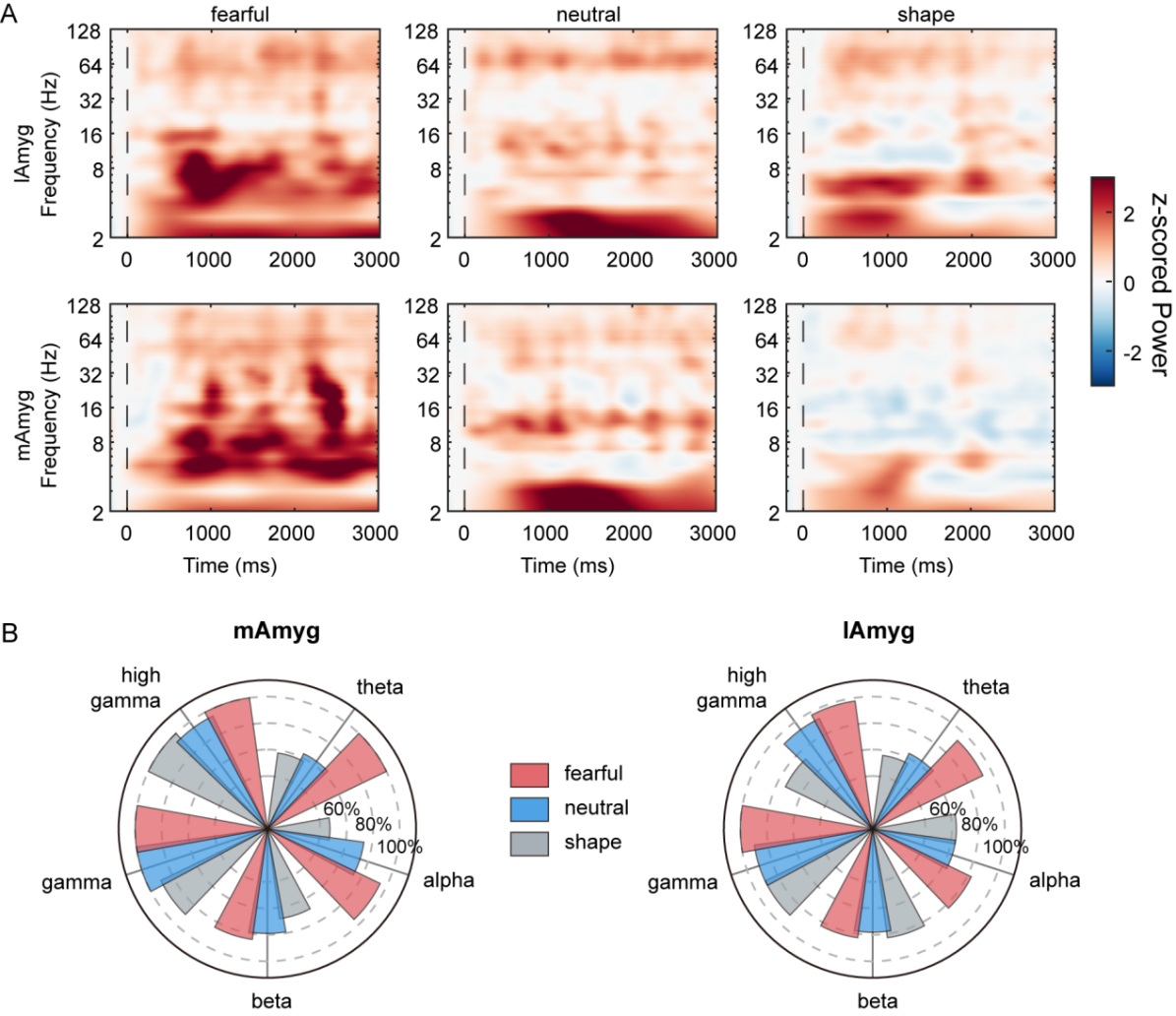

**Fig. S1** Time-frequency power of fearful, neutral and shape conditions across all contacts, and frequency-band distributions of responsive contacts. (A) Group-averaged time-frequency plots of *z*-scored power across all contacts for the fearful (left), neutral (middle) and shape conditions (right) separately. The top row represents the lAmyg, while the bottom row represents the mAmyg. The response contacts serve as the primary source of the observed *z*-scored power across all contacts. (see **Fig. 3A** and **Fig. 5A**). (B) Proportion of responsive contacts across different frequency bands for the fearful (red bar), neutral (blue bar), and shape (gray bar) conditions. The largest proportion of responsive contacts across all frequency bands occurred under the fearful condition (lAmyg: 91.22% ± 8.02%, mAmyg: 95.79% ± 6.86%).

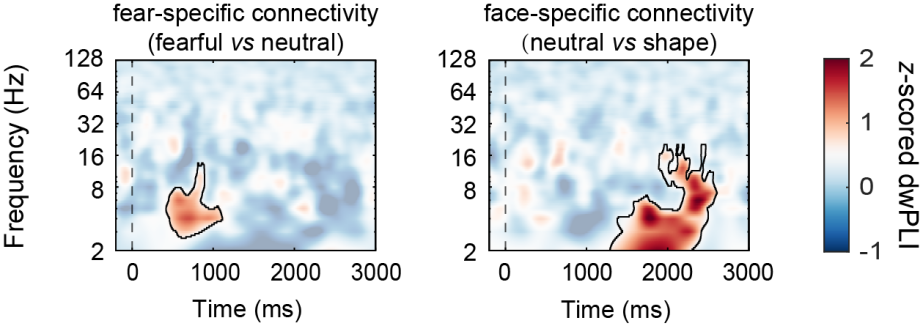

**Fig. S2** Group-level *z*-scored dwPLI between subregions under the fear-specific (left) and face-specific connectivity. Significant phase synchronization was found in the low-frequency range (*z* > 1.96, *ps* < 0.05).
